# Infant Weight Gain Trajectories Linked to Oral Microbiome Composition

**DOI:** 10.1101/208090

**Authors:** Sarah J. C. Craig, Daniel Blankenberg, Alice Carla Luisa Parodi, Ian M. Paul, Leann L. Birch, Jennifer S. Savage, Michele E. Marini, Jennifer L. Stokes, Anton Nekrutenko, Matthew Reimherr, Francesca Chiaromonte, Kateryna D. Makova

**Author notes:** Corresponding authors: Kateryna Makova, Francesca Chiaromonte, Matthew Reimherr.

## Abstract

Gut and oral microbiome perturbations have been observed in obese adults and adolescents. Less is known about how weight gain in early childhood is influenced by gut, and particularly oral, microbiomes. Here we analyze the relationships among weight gain and gut and oral microbiomes in 226 two-year-olds who were followed during the first two years of life, as part of a larger study, with weight and length measured at seven time points. We used these data to identify children with rapid weight gain (a strong risk factor for childhood obesity), and to derive growth curves with novel Functional Data Analysis (FDA) techniques. The children’s oral and gut microbiomes were sampled at the end of the two-year period, and surveyed with 16S sequencing. First, we show that growth curves are associated negatively with diversity and positively with Firmicutes-to-Bacteroidetes ratio of the oral microbiome – a relationship that is also observed in children with rapid (vs. non-rapid) weight gain. We also demonstrate an association between the gut microbiome and child growth, but only when considering the effect of diet on the microbiome. Lastly, we identify several bacterial genera that are associated with child growth patterns. These results suggest that by the age of two, the oral microbiome may have already begun to establish patterns often seen in older obese individuals. They also suggest that the gut microbiome, while strongly influenced by diet, at age two does not harbor obesity signatures many researchers identified in later life stages.

## INTRODUCTION

One in three children in the United States are overweight or obese (Ogden et al. 2016). Many studies have pointed towards a link between this phenotype and the microbiome of the host in both children and adults (reviewed in (Boulangé et al. 2016)), and linked gut microbiome perturbations and obesity in mice. For instance, germ-free mice inoculated with the microbiota of obese mice were shown to develop obesity, whereas their littermates inoculated with the microbiota of lean mice did not (Turnbaugh et al. 2006; Ridaura et al. 2013). In addition, relative to lean mice, obese mice displayed an increase in Firmicutes and a decline in Bacteroidetes (Ley et al. 2005). More recently, similar patterns have been observed in humans; the gut microbiomes of obese adult and adolescent individuals also frequently have low diversity (Bervoets et al. 2013; Ferrer et al. 2013) and an elevated Firmicutes-to-Bacteroidetes (F:B) ratio (Ley et al. 2006; Bervoets et al. 2013; Ferrer et al. 2013). However, these gut microbiome alterations are not universally linked to obesity (Ley 2010; Boulangé et al. 2016; Sze and Schloss 2016), and study results frequently depend on the methods used (e.g. 16S variable region analyzed, sequencing platform, or computational pipeline) and a number of co-factors that can affect gut microbiome – e.g., diet (David et al. 2014; Xu and Knight 2014), exposure to antibiotics (Yallapragada et al. 2015), the use of non-steroid anti-inflammatory drugs (Rogers and Aronoff 2016), host genetics (Blekhman et al. 2015; Davenport 2016; Gomez et al. 2017), and age (Yatsunenko et al. 2012). On the opposite end of the spectrum from obesity, it has also been shown that there are distinct differences in the microbiome of Indian children with a range of nutritional statuses (from healthy to malnourished) (Ghosh et al. 2014; Dinh et al. 2016). Importantly, to our knowledge, no prior study explored the connection between the gut microbiome and the actual temporal trajectories of weight gain in young children, instead of just investigating the binary differences in outcome of growth. Such trajectories vary greatly among children (Smego et al. 2016), and thus may represent a more informative phenotype than a binary outcome (stunted vs. normal, or obese vs. normal) and may increase statistical power in detecting associations between weight gain and microbiome.

In contrast to the many studies concerning the gut microbiome, very few studies explored the oral microbiome and its relationship with weight gain. These latter studies stem from the observed relationship between periodontal disease prevalence and obesity in adults (Al-Zahrani et al. 2003; Kâ et al. 2013). There is a body of literature (Wade 2013; Said et al. 2014) investigating differences in the oral microbiome in relation to periodontal disease (Liu et al. 2012) and dental caries (Serra e Silva Filho et al. 2014; Said et al. 2014; Chen et al. 2015). A few studies directly investigating the relationship between oral microbiome and obesity, again through the lens of oral health, found differences in the oral microbiome composition of obese vs. lean adults and adolescents (Goodson et al. 2009; Zeigler et al. 2012). One such study in adults pointed to the Bacteroidetes species, *Tannerella forsythia*, as having different prevalence in healthy-weight, overweight, and obese groups (Haffajee and Socransky 2009). In adult women it was found that the *Selenomonas noxia*, a Firmicutes species, can predict overweight status with >98% accuracy (Goodson et al. 2009). In adolescents, Zeigler and colleagues documented an increase of both Firmicutes and Bacteroidetes in obese compared to normal-weight individuals, with no significant difference in the abundance between these two phyla (Zeigler et al. 2012).

Findings on how the gut and oral microbiomes are related to weight gain can only be leveraged if we understand what shapes microbiomes starting from early childhood. A handful of studies investigating the early development of the microbiome found that early childhood is a time marked by dramatic microbiome changes (Palmer et al. 2007; Koenig et al. 2011; Vallès et al. 2014; Bäckhed et al. 2015; Chu et al. 2017). A large seeding of the child’s microbiome occurs as a mother-to-child, vertical bacterial transmission during delivery. Infants delivered vaginally have oral and gut microbiomes similar to their mothers’ vaginal microbiomes (Dominguez-Bello et al. 2010; Lif Holgerson et al. 2011; Chu et al. 2017), while infants delivered via Cesarean section have oral and gut microbiomes that resemble their mothers’ skin microbiomes (Madan et al. 2016). Factors that could influence the composition of this influx are the mother’s weight gain (Leng et al. 2015), diabetes (Hu et al. 2013), and smoking during pregnancy (Li et al. 2016). Nonetheless, differences in microbiomes due to delivery mode might be erased by the mounting effects of other factors as early as six weeks after birth (Chu et al. 2017). Diet is one such factor; the gut microbiome differs between breast-and formula-fed infants (Chu et al. 2017), and the first year after birth also comprises other diet transitions affecting gut and oral microbiomes (Holgerson et al. 2013; Vallès et al. 2014). For instance, high-fat and high-carbohydrate diets have been associated with high and low infant gut microbiome F:B ratios, respectively (Ismail et al. 2011). Aside from diet, antibiotics and acid-reducing drugs (proton pump inhibitors and histamine antagonists) can substantially influence the microbiome in relation to the risk of obesity. For instance, exposure to antibiotics in the first two years was associated with higher weight in later childhood (Ajslev et al. 2011; Trasande et al. 2013; Murphy et al. 2014; Yallapragada et al. 2015), but this link has been recently questioned (Gerber et al. 2016; Ianiro et al. 2016). Also, the use of acid-reducing drugs was associated with decreased gut microbiome diversity in adults and premature infants (<34 weeks gestation) (Gupta et al. 2013; Imhann et al. 2015; Jackson et al. 2016). These drugs are widely prescribed, however we are only now beginning to understand their potential effects on the adult microbiome, and know even less about their interplay with a child’s developing microbiome. Early life transitions and their effects on the microbiome are often described as “chaotic” (Koenig et al. 2011) perhaps because they happen over a short period of time, or because there is not a specific order to their succession (Vallès et al. 2014). Nonetheless, within the first several years of life the children’s microbiomes converge on a more adult-like composition (Yatsunenko et al. 2012; Bäckhed et al. 2015; Dinh et al. 2016).

In this study, we investigated oral and gut microbiomes and their links to weight gain in early childhood, using data from 226 mother-child dyads enrolled in the Intervention Nurses Start Infants Growing on Healthy Trajectories (INSIGHT) study (Paul et al. 2014). Each child’s gut and oral microbiomes were assessed once, at two years of age, and the maternal oral microbiome was assessed at the time of her child’s two-year visit. In contrast, each child’s weight and length were measured at seven time points during the first two years, and we quantitated weight gain with two different approaches: we (a) derived *growth curves* subjected to Functional Data Analysis (FDA) techniques, which exploit mathematical functions formed by the temporal nature of the measurements (Ramsay and Silverman 2007, 2005), and (b) calculated *conditional weight gain* (CWG) *scores*, a traditional metric used to define rapid weight gain in infants (Griffiths et al. 2009; Savage et al. 2016). Both quantitations incorporate information about weight changes during the initial period after birth, but CWG scores only use differences between two time points, while growth curves use several time points longitudinally to fully capture weight dynamics. INSIGHT also includes a broad array of clinical, anthropometric, demographic, and behavioral variables on mother-child dyads (Paul et al. 2014) which we used in our analyses. In detail, we: (a) modeled growth curves and examined their associations with oral and gut microbiomes; (b) assessed whether oral and gut microbiomes differ between children with rapid vs. non-rapid weight gain; (c) investigated whether there is any relationship between the maternal oral microbiome and her child’s weight trajectory, and (d) analyzed whether diet and other factors have an impact on oral and gut microbiomes and/or the growth of the child. We used a variety of statistical tools, including FDA methods recently developed by our group, and developed a freely accessible pipeline for analyzing 16S rRNA gene sequences on Galaxy (usegalaxy.org). Our results demonstrate significant associations between the oral microbiome, as established at age two, and rapid weight gain during the first two years of life. They also capture significant effects of diet on the gut microbiome at age two that may modulate specific growth trajectories in children.

## RESULTS

### Oral and gut microbiome profiling

From the 279 families recruited in INSIGHT that had complete longitudinal anthropometric and behavioral measurements over the first two years after birth (Paul et al. 2014), we collected samples from 226 mother-child dyads (specifically, oral samples for 215 mothers and 214 children, and stool samples for 189 children; see Fig. S1 for a graphical summary) at the child’s two-year clinical research lab visit. For these samples, we sequenced the variable regions 3 and 4 of the 16S rRNA gene and analyzed the data in Galaxy (Afgan et al. 2016) to determine taxonomic classifications and compute ecological diversity measures (see Methods).

Overall, our oral and gut microbiome samples showed fundamental differences in composition at the phylum level; oral microbiomes harbored abundant Fusobacteria and no Verrucomicrobia, whereas the gut microbiomes harbored abundant Verrucomicrobia but no Fusobacteria (Figs. S2A-B). These results are largely consistent with previously published studies exploring the composition of healthy, adult gut (Lozupone et al. 2012) and oral (Dewhirst et al. 2010) microbiomes. Qualitatively, it appears that a child’s oral microbiome (Fig. S2A) is more similar to his/her mother’s oral microbiome (Fig. S2C) than to the child’s gut microbiome (Fig, S2B). Additionally, within each microbiome type (child oral, child gut, and maternal oral) the proportions of bacteria belonging to different phyla varied greatly among individuals (Figs. S2A-C). Using non-metric multidimensional scaling (NMDS) to compare the three microbiome types (Fig. S2D) we found that there was some overlap between oral and gut microbiome communities, but stool samples largely clustered together and away from oral samples (mother and child), and children’s oral samples formed a tighter cluster than maternal oral samples. These results suggest a higher interindividual variability with age, as well as the development of distinct microbiome communities at different body sites by the age of two years.

### Growth curves and their relationship with the microbiome

Children’s growth curves were constructed based on the ratio between weight and length (later called *growth index*) measured at seven time points during the first two years after birth (see Methods). This set of longitudinal observations (Fig. 1A and Fig. S3) was processed with FDA techniques (Yao et al. 2005): the growth indices of all children were pooled to estimate population-level curve parameters (mean and covariance functions), and smooth individual growth curves were constructed forming best linear unbiased predictors of the missing segments (see Methods). These curves were monotonously increasing (Fig. S3B), highlighting fast growth rate over the first two years after birth. Because each child may hit growth spurts at a slightly different age (Smego et al. 2016), we aligned the curves based on their dominant shapes (Fig. 1B). The alignment procedure had only minor effects on the curves (compare Fig. S2B and Fig. 1B) but allowed us to focus on variation in the amplitude of the curves (i.e. growth index on the vertical axis), while reducing variation in their temporal phasing (i.e. time on the horizontal axis).

**Fig. 1.**
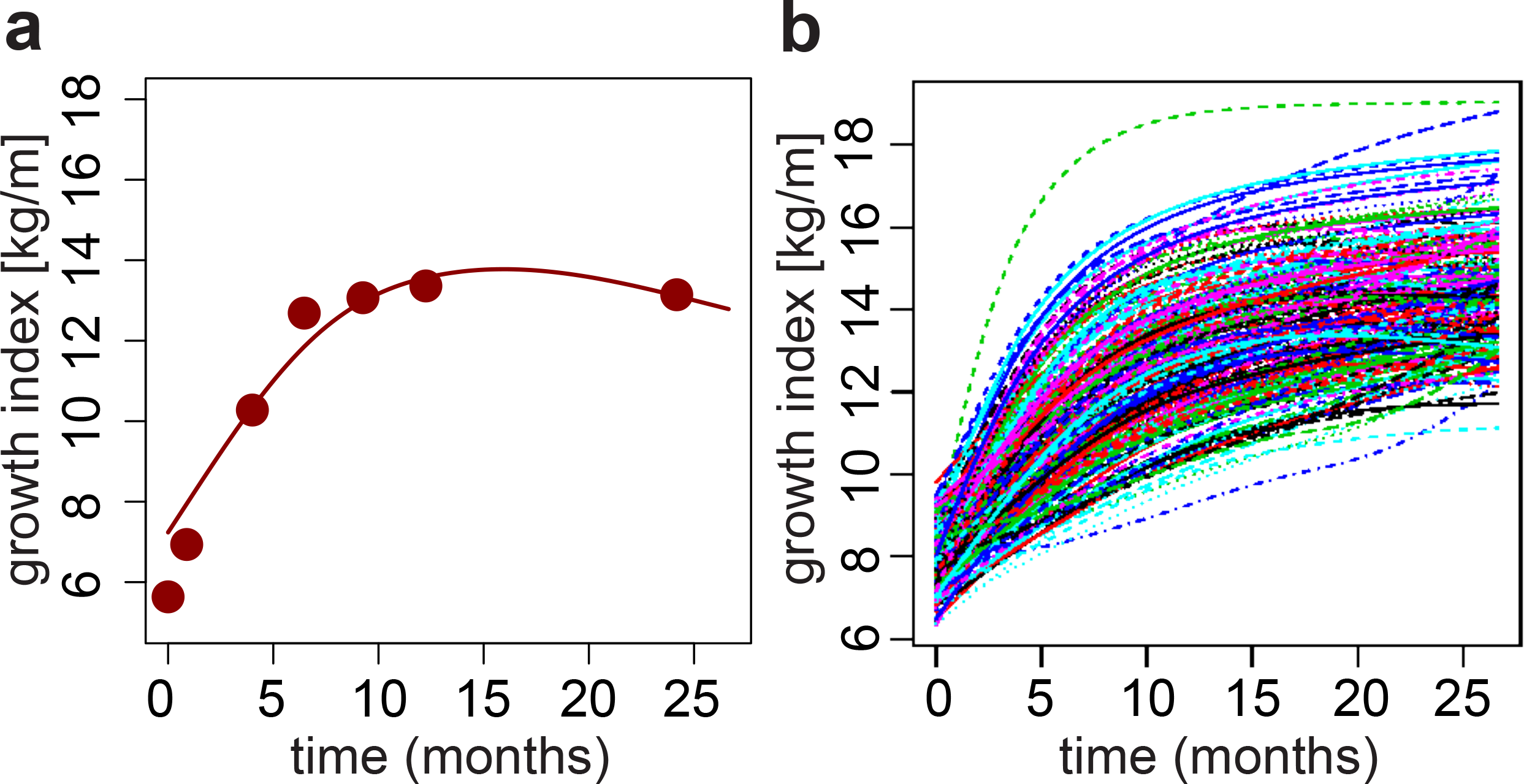
Growth curves construction. (**A**) Example growth curve. Points: observed weight-to-length ratios (i.e. growth indices); line: estimated growth curve. (**B**) Final aligned growth curves for all children studied.

To evaluate associations between children’s weight gain and the microbiome information, we started by applying FDA regression techniques (see Methods). Using children’s growth curves (Fig. 1B) as the response we ran a total of six functional regressions, each with a single scalar predictor – children’s oral α-diversity, oral F:B ratio, gut α-diversity and gut F:B ratio; and mothers’ oral α-diversity and oral F:B ratio. In these functional regressions, one has regression *coefficient curves* – as opposed to scalar regression coefficients. Estimated coefficient curves above/below the zero line suggest negative and positive associations. Significance of associations can be measured through p-values obtained from various tests (see Table S1) and also gleaned by whether the zero line is contained within a curve’s 95% confidence band. Our results suggest that low microbial diversity and high content of Firmicutes relative to Bacteroidetes in the mouth of a two-year-old are markers of elevated growth indices during the first two years after birth – the estimated coefficient curves are negative and positive, respectively, with confidence bands that do not contain the zero line (Figs. 2B and 2E), and p-values are below 5% (Table S1). Conversely, estimated coefficient curves for the gut are closer to the zero line (Figs. 2A and 2D) and p-values are large (Table S1), suggesting that α-diversity and F:B ratio in the gut of a two-year-old are not significantly associated with his/her growth trajectory from birth to two years. Interestingly, α-diversity, but not the F:B ratio, in mothers’ oral microbiomes are significantly associated with their children’s growth curves (Figs. 2C and 2F, and Table S1). In fact, the α-diversities of children’s and mothers’ oral microbiomes were significantly correlated (R = 0.20, p = 0.003), and the corresponding estimated regression coefficient curves had similar shapes (Figs. 2B-C).

**Fig. 2.**
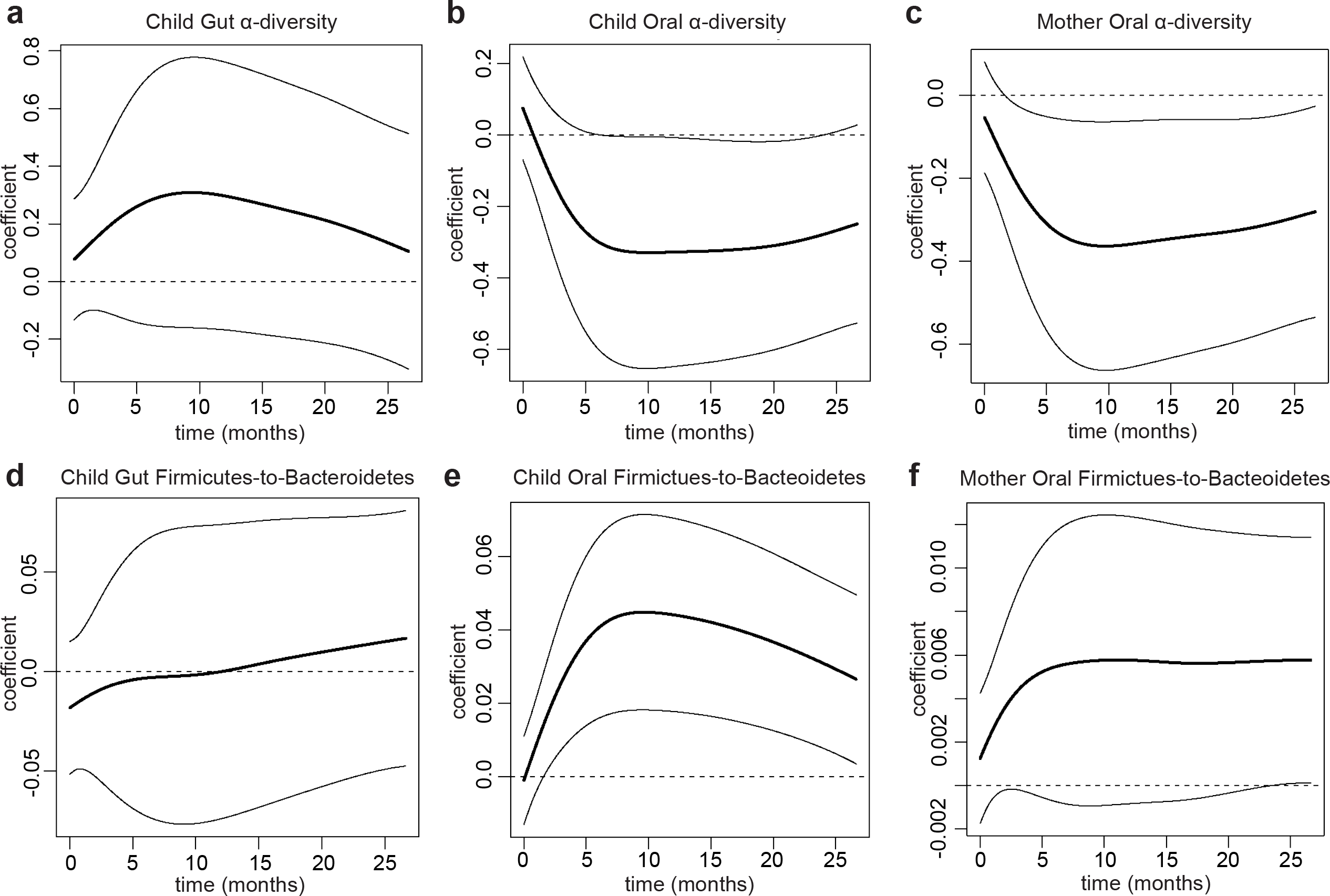
Oral and Gut microbiome’s relationships with growth curves. Estimated regression coefficient curves expressing the associations of growth curves with (**A**) children’s gut α-diversity, (**B**) children’s oral α-diversity, (**C**) mothers’ oral α-diversity, (**D**) children’s gut Firmicutes-to-Bacteroidetes ratio, (**E**) children’s oral Firmicutes-to-Bacteroidetes ratio and (**F**) mothers’ oral Firmicutes-to-Bacteroidetes ratio. Each curve is accompanied by a point-wise confidence band (Ramsay and Silverman 2005).

### Rapid infant weight gain and its relationship with the microbiome

To complement the analyses based on growth curves, we associated the binary quantification of rapid vs. non-rapid weight gain, as defined by Conditional Weight Gain (CWG) scores, to the microbiome information. Children’s CWG scores were computed from weight gain between birth and six months, corrected by length at these two time points (Griffiths et al. 2009; Savage et al. 2016) (see Methods). A CWG score ≥ 0 indicates weight gain that is faster than average and is used to define rapid weight gain (Savage et al. 2016) – a predictor of obesity later in life (Taveras et al. 2009). In our cohort, 103 children had rapid weight gain between birth and six months (CWG ≥ 0) and 123 did not (CWG < 0). Notably, the former had a significantly greater weight at two years of age than the latter (p=4.627e-13, Mann-Whitney one-tailed t-test; Fig. S4).

To test for associations between CWG scores and microbiomes, first, we assessed whether the microbiome of children with rapid weight gain possessed the obesity signatures previously found in the gut microbiome of older obese children and adults (see Introduction) – namely, lower diversity and higher F:B ratio than non-rapid weight gain children. This was indeed the case for the oral microbiome (p = 0.049 and p = 0.019, respectively, one-tailed Mann-Whitney U test; Figs. 3B and 3E) but, interestingly, not for the gut microbiome (p = 0.776 and p = 0.286, respectively; Figs. 3A and 3D). Second, we tested whether mothers of children with rapid vs. non-rapid weight gain had significantly different oral microbiome -diversities and F:B ratios. Again we found that diversity was significantly lower in mothers of children with rapid weight gain (p = 0.036, Fig. 3C), but the F:B ratio was not significantly higher (p = 0.093, Fig. 3F). The patterns in these analyses are consistent with those in the growth curves analyses, but the latter provided stronger significance assessments (Table S1), demonstrating the effectiveness of FDA techniques – which incorporate longitudinal information in a richer and more nuanced fashion.

**Fig. 3.**
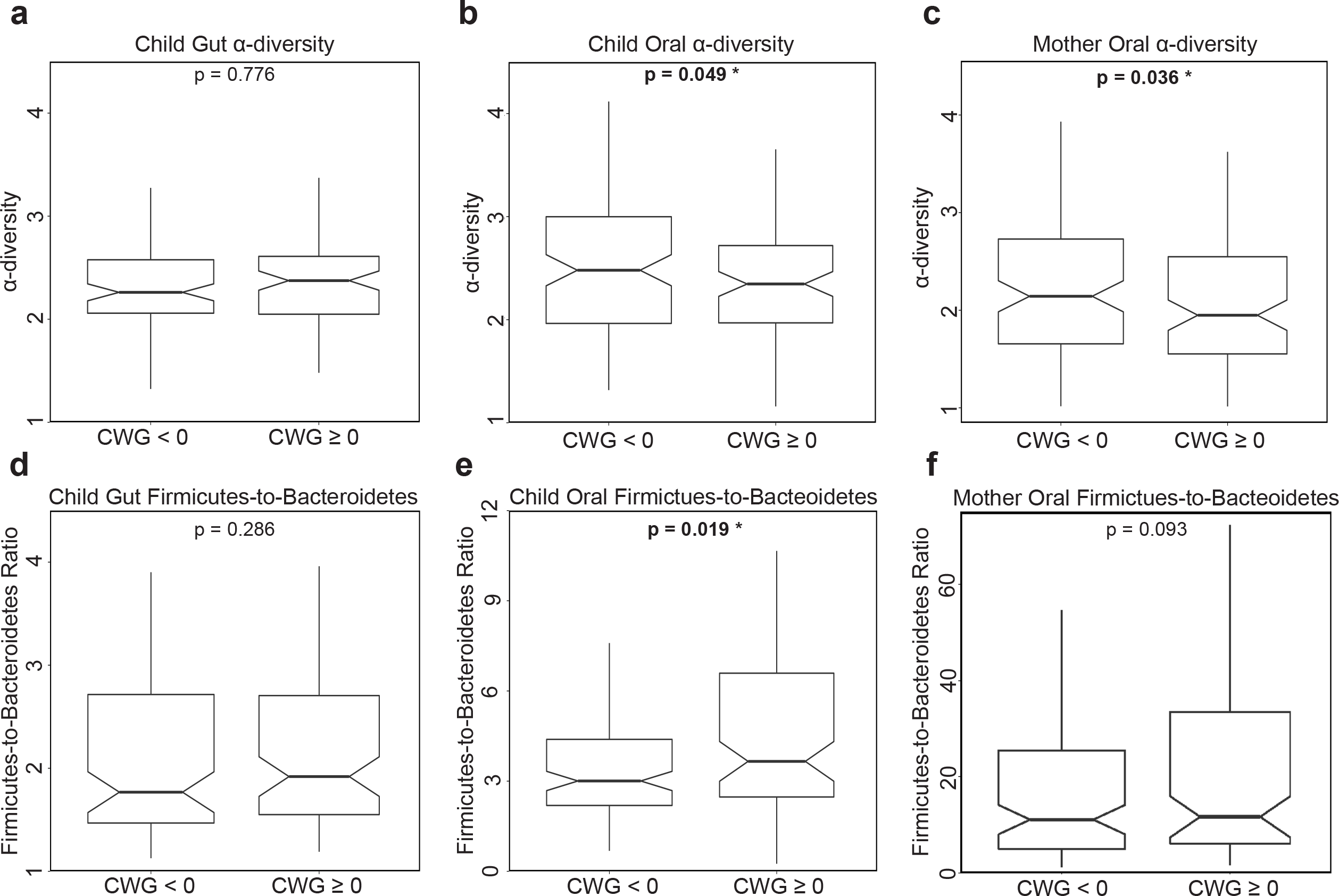
Oral and Gut microbiome’s relationships with conditional weight gain. Notched box-plots contrasting α-diversity and Firmicutes-to-Bacteroidetes (F:B) ratio in two-year-old children with rapid (CWG ≥ 0) vs. non-rapid (CWG < 0) weight gain, and in their mothers. (**A, D**) for the *gut* microbiome in children with rapid (N = 90) vs. non-rapid (N = 99) weight gain; (**B, E**) for the *oral* microbiome in children with rapid (N = 97) vs. non-rapid weight gain (N = 117); (**C, F**) for the oral microbiome in mothers of children with rapid weight gain (N = 102) vs. mothers of children without rapid weight gain (N = 113). All p-values were obtained using one-tailed Mann-Whitney U tests. Outliers were not plotted but were included in the tests.

### Potential co-factors and their roles

Many factors could affect children’s microbiomes and their relationships with weight gain. Based on the anthropometric and behavioral data at our disposal (Paul et al. 2014), we considered gender, exposure to antibiotics and acid-reducing medication during the first two years after birth, delivery mode (vaginal vs. C-section), INSIGHT intervention (Paul et al. 2014), maternal gestational diabetes, gestational weight gain, smoking during pregnancy, family income, and diet information (see Methods). These factors were tested, both individually and jointly through multiple regressions, for effects on our four summary measures of children’s microbiomes; namely, children’s oral and gut α-diversity and F:B ratio at two years of age. Results of these tests were largely non-significant, possibly due to the small number of subjects in some of the conditions considered (Table S2). However, diet at age two emerged as having marked effects on the gut microbiome at the same age. Multiple regressions on diet-related variables showed strong effects on both gut F:B ratio and α-diversity (Table S3). We also ran variable selection for multiple regressions comprising seventeen potential covariates (from Tables S2 and S3). Notably, only diet-related variables were selected by the procedure, and only in regressions for the gut microbiome (no covariates were retained in regressions for the oral microbiome). Fitting regressions restricted to the selected diet-related variables, we explained as much as 20.79% of the variability in gut F:B ratio, with vegetable and meat consumption as significant positive and negative predictors, respectively, and a more modest 5.94% of the variability in gut α-diversity, with vegetable and fruit consumption as significant negative and positive predictors, respectively (Table 1).

**Table 1.**
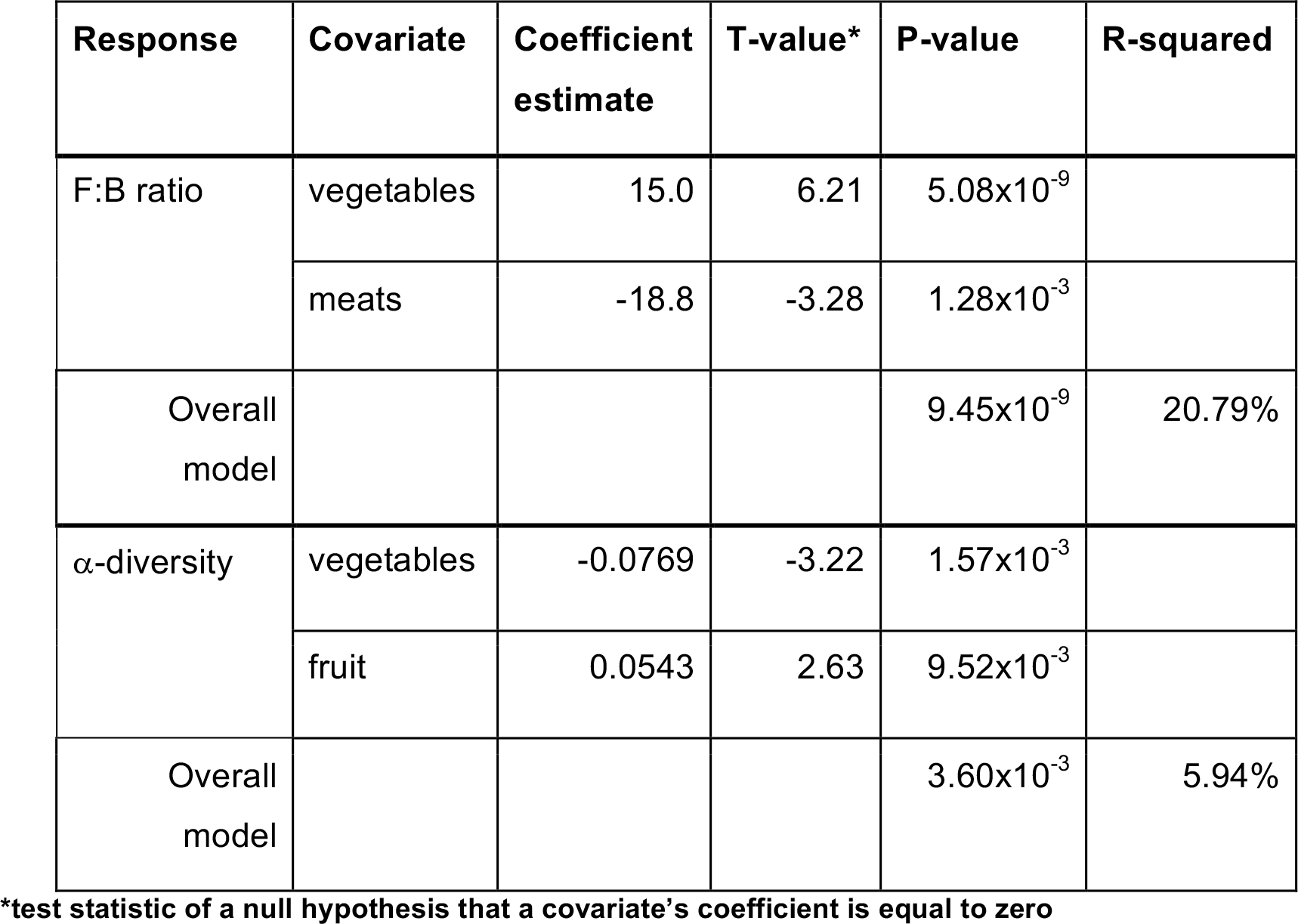
Associations detected in multiple linear regressions for children’s gut microbiomes’ Firmicutes-to-Bacteroidetes ratio and α-diversity. Each regression comprised 17 potential covariates, but only diet-related ones were retained by variable selection procedures. Coefficient estimates and significance (p-values) are shown, together with overall model significance and R-squared, for the regression restricted to the selected diet-related covariates.

| Response | Covariate | Coefficient estimate | T-value* | P-value | R-squared |
| --- | --- | --- | --- | --- | --- |
| F:B ratio | vegetables | 15.0 | 6.21 | $5.08 \times 10^{-9}$ | |
| | meats | -18.8 | -3.28 | $1.28 \times 10^{-3}$ | |
| Overall model | | | | $9.45 \times 10^{-9}$ | 20.79% |
| α-diversity | vegetables | -0.0769 | -3.22 | $1.57 \times 10^{-3}$ | |
| | fruit | 0.0543 | 2.63 | $9.52 \times 10^{-3}$ | |
| Overall model | | | | $3.60 \times 10^{-3}$ | 5.94% |

\*test statistic of a null hypothesis that a covariate's coefficient is equal to zero

Interestingly, functional regressions of our growth curves on diet-related variables did not indicate any significant effects (Table S4). Nevertheless, because of its strong effects on the gut microbiome at age two, diet may be a modulator of its relationships with weight gain. To assess this, we repeated the functional regressions of growth curves on gut microbiome summary measurements adding diet-related covariates with significant effects on the microbiome (from the previous variable selection procedure, see Table 1). Notably, in one of the statistical tests we ran, gut α-diversity became a significant positive predictor of growth curves when considered together with diet (Table S5 and Fig. S5A), while gut F:B remained non-significant in all our tests (Table S6 and Fig. S5B).

### Weight gain and microbiome composition: influential taxa

Next, we went beyond α-diversity and F:B ratio summary measures, which were computed at the coarse phylum level, and considered microbiome composition at a finer resolution to identify bacterial genera associated with a child’s weight gain. We computed *genus-level*, normalized abundances for the same 214 children’s oral microbiomes, 189 children’s gut microbiomes, and 215 mothers’ oral microbiomes used for the analyses in the previous sections. Because many of these abundances were very low or highly collinear, we implemented a procedure to aggregate them into *taxonomic groups* (below we refer to these also as *bacterial groups*, or simply *groups*), leveraging the phylogeny of bacterial genera (see Methods; about 25% of the groups comprised just one bacterial genus, but 75% aggregated two or more genera that were either very scarce or highly correlated). We obtained 75, 77, and 79 taxonomic groups for children’s oral, children’s gut, and mothers’ oral microbiomes, respectively (Table S7). Based on the resulting aggregated abundances, and separately for the three microbiomes, we used FLAME (a novel FDA methodology recently developed by our group (Choi and Reimherr 2017; Parodi and Reimherr 2017)) to identify taxonomic groups that are the best predictors of children’s growth curves (Table S8).

In the children’s gut microbiome FLAME detected a group from the Proteobacteria phylum with a negative effect (Group 61), three Firmicutes groups with positive effects (Groups 23, 26, and 41) and one Bacteroidetes group with a positive effect (Group 12; Fig. 4A). In the children’s oral microbiome FLAME detected a group of Bacteroidetes (Group 13) and a group of Actinobacteria (Group 5) having, respectively, positive and negative effects (Fig. 4B). Two groups were detected in the mothers’ oral microbiome (Fig. 4C) – one Firmicutes group with a positive effect (Group 28) and one Fusobacteria group with a negative effect (Group 53).

**Fig. 4.**
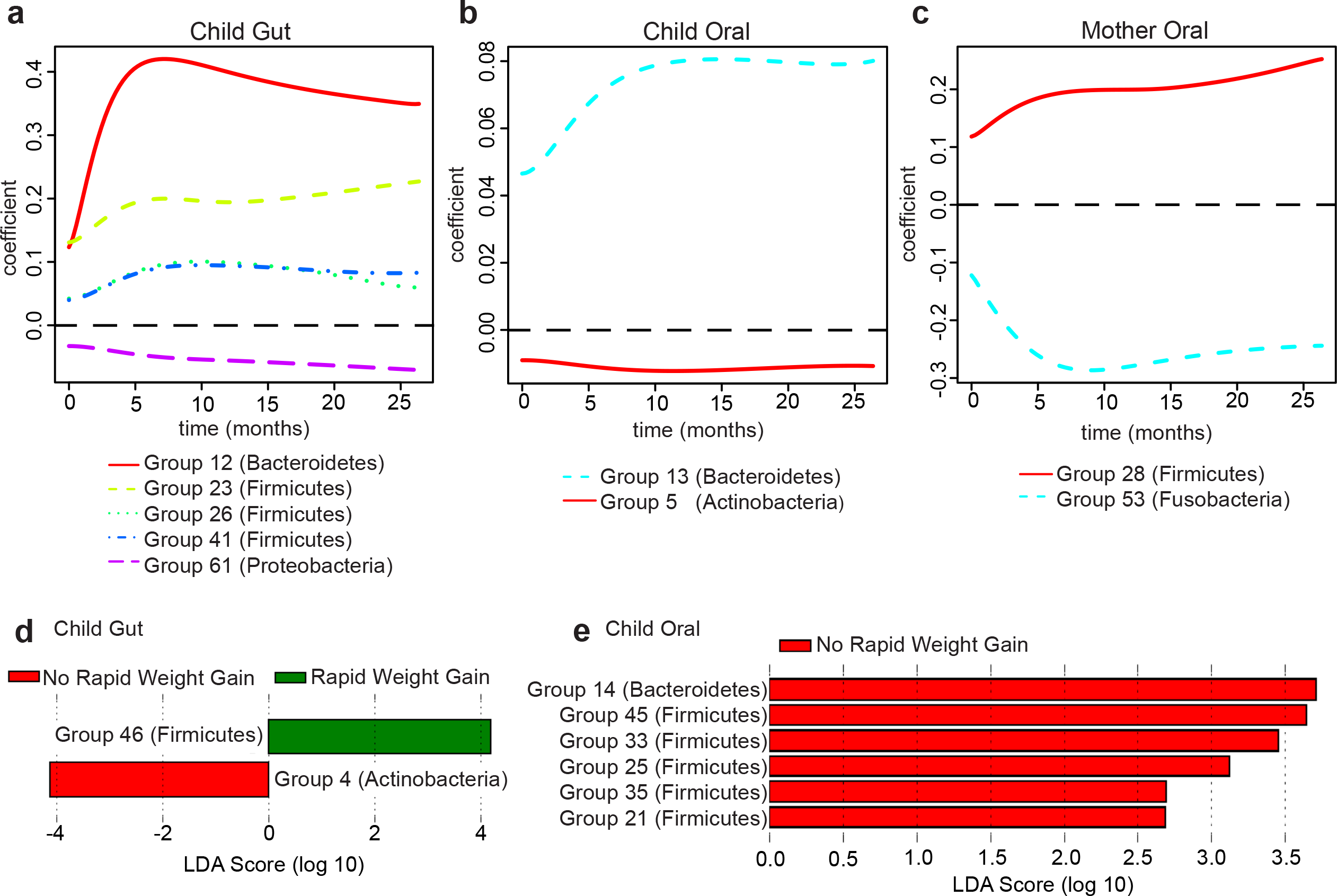
Identification of influential taxonomic groups. Estimated regression coefficient curves for taxonomic groups affecting growth curves, as identified by FLAME (Parodi and Reimherr 2017) in: (**A**) children’s gut microbiome, (**B**) children’s oral microbiome, (**C**) mothers’ oral microbiome. The zero line (dashed) corresponds to no effect. Linear Discriminant Analysis scores for taxonomic groups distinguishing children with rapid vs. non-rapid weight gain, as identified by LEfSe (Segata et al. 2011) in: (**D**) children’s gut microbiome, (**E**) children’s oral microbiome. See Table S8 for a list of genera belonging to each group identified by FLAME and LEfSe.

We also used the Linear Discriminant Analysis (LDA) effect size tool, LEfSe (Segata et al. 2011), to identify taxonomic groups that were most informative for separating children with rapid vs. non-rapid weight gain (Table S8). In the children’s gut microbiomes, LEfSe detected two discriminant bacterial groups; an Actinobacteria group associated with non-rapid weight gain and a Firmicutes group associated with rapid weight gain (Groups 4 and 46; Fig. 4D). In the children’s oral microbiomes, LEfSe detected several discriminant groups associated with non-rapid weight gain, belonging to the Bacteroidetes and Firmicutes phyla (Groups 14, 21, 25, 33, 35, and 45; Fig. 4E). LEfSe did not identify any discriminant groups in the mothers’ oral microbiome.

## DISCUSSION

### Children’s oral microbiome and weight gain

Our results demonstrate that a child’s *oral* microbiome, as analyzed at age two, bears significant associations with weight gain during the first two years after birth. In particular, it displays the decreased diversity and increased F:B ratio signatures characteristic of the *gut* microbiome of obese adolescents and adults (Bervoets et al. 2013). This conclusion was supported by sophisticated FDA techniques leveraging longitudinal information on weight gain in the first two years after birth, and confirmed by standard statistical tests based on a binary phenotype (rapid vs. non-rapid weight gain). Notably, links between the oral microbiome and a child’s weight gain trajectory have not been previously demonstrated for this age group, nor for a healthy population. Altogether, our results suggest that the association between the oral microbiome and the temporal pattern of weight gain in early childhood might be stronger and more consequential than previously thought, and thus requires further characterization.

Why does the oral microbiome carry markers of rapid weight gain in children? Some studies have pointed to potential mechanisms linking periodontal disease and obesity, including increased oxidative stress (Bullon et al. 2009), low-grade systemic inflammation and insulin resistance (Li et al 2009, Kwon et al 2011), and higher gingival crevicular fluid TNF- (Lundin et al. 2004, Kâ et al. 2013). Goodson and colleagues (Goodson et al. 2009) suggested that the oral microbiome could (a) affect the gastrointestinal tract to increase metabolic efficiency, resulting in increased fat storage; (b) affect leptin or ghrelin levels, resulting in increased appetite and food consumption; and/or (c) affect TNF- and adiponectin pathways, resulting in insulin resistance and increased fat storage. Gingival inflammation and decrease in salivary secretion rate were also observed in children with obesity, without distinct microbial profiles when compared to normal weight peers (Fadel et al. 2013). Deciphering causal mechanisms through which the oral microbiome might affect weight gain is outside the scope of this study, especially since we linked microbiomes assayed at age two with growth trajectories prior to that age. However, the associations we detected should be studied further using data collected longitudinally not just for body size, but also for the microbiome. If confirmed in such a study and for other populations, the relationships we gleaned between the oral microbiome and weight gain could lead to a non-invasive clinical screen to identify children who are at a particular risk of developing obesity later in life. These at-risk children could be closely monitored and be the primary candidates for obesity-prevention interventions (Paul et al. 2014).

Notably, we also detected a significant association between mothers’ oral microbiome diversity and their children’s growth curves, suggesting the presence of a familial (genetic and/or household) microbiome signature linked to the dynamics of weight gain in early childhood. This too should be explored further, collecting and analyzing longitudinal data on the oral microbiomes of parents and children.

### Children’s gut microbiome and weight gain

In contrast to the oral microbiome, we find that a child’s gut microbiome at age two is not significantly associated with weight gain during the first two years after birth. At first, this appears surprising, as several studies have linked obesity to decreased diversity and increased F:B ratio in the gut microbiome. However, while an increased F:B ratio is a common marker of obese gut microbiomes described in several papers (Turnbaugh et al. 2006; Ley et al. 2006) and reviews (Ley 2010; Boulangé et al. 2016), some studies found no change in Bacteroidetes, increased Bacteroidetes in overweight and lean individuals, or increased Firmicutes in lean patients after gastric bypass (reviewed in (Ley 2010; Boulangé et al. 2016)). Moreover, the fact that we are studying a very young and not yet completely established gut microbiome might at least partially explain the absence of signal in our data; the obesity signatures of decreased diversity and increased F:B ratio may become pronounced only at later stages of gut microbiome development. This may suggest that the oral microbiome is established with potential signatures of obesity earlier than the gut microbiome. We also note that, despite the lack of significant associations between gut microbiome summary measures and growth curves (or binary weight gain outcome), we did find specific gut taxonomic groups associated with early childhood weight gain (see below). These could be pioneers in setting up changes leading to decreased diversity and increased F:B ratio – a hypothesis that should be investigated in future studies.

### Diet, microbiomes and weight gain in children

We had data on a large number of factors potentially affecting a child’s gut and oral microbiomes, but with the notable exception of diet-related variables for which we found strong effects on the gut microbiome, their effects were non-significant or inconclusive. This could have been due, at least partially, to the small number of subjects in some of the conditions considered. Another potential explanation is that the microbiomes were characterized at the age of two, while most factors (except for current diet information) were measured at earlier time points – so their effects might have diluted over time. Sampling the microbiomes at multiple time points during early childhood should alleviate this limitation in future studies and will allow us to further exploit FDA to analyze longitudinal information not only on growth curves, but also on microbiome development.

Interestingly, diet-related variables at the age of two did *not* show significant associations with weight gain between birth and two-years (no significant association was detected when regressing growth curves on gut microbiome α-diversity or F:B ratio; Table S4). However, perhaps because of the diet variables’ strong effects on the gut microbiome (Table 1), they appear to modulate the relationship between weight gain and the gut microbiome – when adding diet-related variables in a joint regression with the microbiome, α-diversity became a significant positive predictor of child growth trajectories (F:B ratio remained non-significant). In other words, if we compared the gut microbiome of children with the same diet, children who gain weight more rapidly would harbor greater microbial diversity. Thus, a diversity signature in the gut microbiome appears to emerge in our data when controlling for diet, albeit with a sign opposite to the one most commonly identified in the literature – and also opposite to the one we detected in the oral microbiome. We presently do not have an explanation for this finding.

### Influential taxa and the power of novel methodology

It is surprising at first sight that, while no significant (either positive or negative) association was found between gut microbiome F:B ratio and weight gain, both Firmicutes and Bacteroidetes were identified as influential groups by our FLAME and/or LEfSe analyses. However, diet might have influenced these patterns. For instance, FLAME identified a Bacteroidetes group in the gut as positively associated with children’s growth curves, and using linear regression we found that meat consumption has a negative effect on gut F:B ratio (which could be due to a positive association with Bacteroidetes in the denominator). Additionally, meat consumption was significant in the functional regression of growth curves on gut F:B ratio and diet-related variables (even though F:B ratio itself was not). Thus, while other researchers have found that individuals with a “Western diet” (high in animal protein) have increased gut Bacteroidetes abundances (Wu et al. 2011), the exact nature of these relationships cannot be completely elucidated with the data currently at our disposal.

Another somewhat surprising finding is that the taxonomic groups identified by FLAME and LEfSe do differ. This could potentially be due to inherent methodological differences between FLAME (a tool that selects groups using high-dimensional function-on-scalar regressions) and LEfSe (a tool using Linear Discriminant Analysis on differentially abundant groups). Alternatively, these incongruencies could be due to the very different ways in which the weight gain phenotype is encoded (binary outcome vs. temporal change); and therefore it may be a biological explanation of why bacteria influencing the shape of the growth curves might in fact differ from those discriminating between the microbiomes of children with rapid vs. non-rapid weight gain. This will require further examination in future studies, as our sequencing data currently do not allow resolution at the species or strain level, and the functional capacity of the microbiome is unknown. Furthermore, sampling the microbiome longitudinally, along with biometric measurements (instead of at the single two-year time point available to us in this study), may unveil additional bacterial groups relevant for the perspective (instead of retrospective) relationship with weight gain patterns. Similar considerations apply to the fact that, with the data at our disposal, FLAME identified different bacterial groups in children’s and mothers’ oral microbiomes (Fig. 4B-C).

Nonetheless, the use of FDA techniques afforded us the opportunity to associate longitudinal information (growth curves derived from weight and length collected at multiple time points during the first two years after birth) with microbiome measures at the age of two, which, to our knowledge, has not been done before. We used FDA in addition to more traditional statistical analyses based on a binary phenotype (rapid vs. non-rapid weight gain). While results on the relationship between growth and microbiome diversity and F:B ratio were consistent between the two approaches, growth curves captured weight gain in a richer fashion than the binary phenotype – leading to stronger results significance. Growth curves also led to additional insights. For instance, a larger number of taxonomic groups were found to be significantly associated with growth curves using FLAME than with rapid vs. non-rapid weight gain phenotypes using LEfSe. Our study demonstrates the effectiveness of FDA techniques linking children’s weight gain trajectories and microbiomes characterization. This suggests that even greater effectiveness could be achieved in similar studies with longitudinal information also on the microbiomes, and indicates the potential of such techniques in a variety of other ‘omics’ applications (e.g.,(Campos-Sánchez et al. 2016)).

## METHODS

### Study population

We collected microbiome information on 226 mother-child dyads recruited from the 279 families involved in the INSIGHT study (Paul et al. 2014). These dyads included full-term singletons born to primiparous mothers in Central Pennsylvania, and were predominantly white (Savage et al. 2016). The INSIGHT study collected clinical, anthropometric, demographic and behavioral variables on children and mothers (Paul et al. 2014). Parents completed questionnaires reporting children’s dietary intake and exposure to medications. Children’s weight and length were measured at birth, 3-4 weeks, 16 weeks, 28 weeks, 40 weeks, 1 year, and 2 years.

### Microbiome collection and sequencing

Buccal samples were collected by research staff of the Penn State Hershey Pediatric Clinical Research Office at the child’s two-year clinical research center visit. Ten sterile cotton swabs were each rubbed for 20 seconds against the inside cheeks of children or mothers. The cotton swabs were placed in tubes containing slagboom buffer (Freeman et al. 2003). Samples were stored in the Pediatric Clinical Research Office and transported from the Hershey Medical Campus to the University Park Campus where they were processed.

Right before the two-year visit, stool samples were collected by parents in stool collection tubes, wrapped in freezer packs, and frozen immediately in the home freezer. They were then brought packed on ice to the clinical research site by the parents and stored at −20°C. Samples were finally transported in coolers on ice from the Hershey Medical Campus to the University Park Campus where they were stored at −80°C until processing. DNA was extracted using the MoBio PowerSoil DNA Isolation kit. The variable regions 3 and 4 of the 16S rRNA gene were amplified and sequenced on an Illumina MiSeq (v3 chemistry, 2x300 reads).

### Sample DNA extraction, library preparation, and DNA sequencing

Genomic DNA (gDNA) was extracted from samples using the MoBio PowerSoil DNA isolation kit (Qiagen). The manufacturer’s directions were followed with modifications implemented based on the protocol established by the Human Microbiome Project. These include two heating steps (65°C and 90°C for 10 minutes each) prior to bead beating. To control for contamination in the DNA from this stage, a blank ‘sample’ (containing only MoBio Bead Buffer) was subjected to the entire DNA extraction protocol and library preparation. Contamination was never detected via gel electrophoresis, Qubit, or Bioanalyzer.

Library preparation for sequencing of the 16S rRNA gene followed the Illumina protocol ‘16S Metagenomic Sequencing Library Preparation’ (Illumina part# 15044223 Rev.B; https://support.illumina.com/downloads/16s_metagenomic_sequencing_library_preparation.html). Each library was quantified using a fluorometer (Qubit, ThermoFisher Scientific) and analyzed for correct size on a BioAnalyzer (Agilent). As a positive control, a synthetic mock community DNA pool (BEI Resources) was amplified and sequenced alongside the experimental samples. To control for contamination in the amplification or library preparation steps, a blank sample was added to all of the library preparation steps. Contamination was never detected via gel electrophoresis, Qubit, or Bioanalyzer.

An equimolar pool of 48 libraries with a 25% PhiX spike-in was sequenced on an Illumina MiSeq (v3 chemistry, 2 x 300 reads). Demultiplexing was performed using the Illumina software. FASTQ files were retrieved to be used in downstream analyses.

### Sequence analysis and diversity measures

Sequences were first examined using FastQC (Andrews 2010) and a multi-sample report was generated for textual and graphical views using Galaxy tools (Blankenberg et al. 2010a) and MultiQC (Ewels et al. 2016), respectively. FASTQ manipulation filters (Blankenberg et al. 2010a) were then applied to remove low-quality sequences. Sequences were trimmed using a sliding window approach. To both the 5’ and 3’ ends of the sequences, we applied a cutoff of minimal mean PHRED score of 20 within a window of 5 bases, step size of 1. Quality was reassessed after trimming.

We then merged/joined our forward and reverse paired-end reads into a single contig using a two-step process. First, we used fastq-join from the *ea-utils* package (Aronesty 2011; Aronesty, 2013) to merge overlapping forward and reverse reads into a single read. This process aligns read pairs and merges overlapping regions based upon user-specified parameters of mismatch percentage and minimum alignment length; we utilized 8% and 6 bases, respectively. To prevent the loss of non-mergeable reads, we performed a second joining operation, using the FASTQ Joiner tool (Andrews 2010; Blankenberg et al. 2010) and inserted a string of 5 ambiguous nucleotides (“NNNNN”) between the pairs. We next removed chimeric sequences. We used VSearch (Rognes et al. 2016) with the uchime_denovo algorithm to create a list of non-chimeric sequences.

Non-chimeric reads were classified individually into taxa. We utilized Kraken (Wood and Salzberg 2014) with a customized database containing only 16S rRNA gene sequences, based on GreenGenes (DeSantis et al. 2006). Once Kraken had assigned taxa-kmer counts to individual reads, we utilized a custom abundance reporting tool (https://github.com/blankenberg/Kraken-Taxonomy-Report) to report abundances across samples at specified ranks, along with a phylogenetic tree that was pruned to contain the terminal nodes that are present in at least one of the samples and the connected internal nodes.

Rarefaction and α- and β-diversity calculations were performed using Vegan (Jari Oksanen, F. Fuillaume Blanchet, Roeland Kindt, Pierre Legendre, Peter R. Minchin, R. B. O’Hara, Gavin L. Simpson, Peter Solymos, M. Henry, H. Stevens, Helene Wagner 2015). Rarefaction was used to normalize the read count across samples – which were all rarefied to 100,000 reads. To compute summary measures of diversity for each microbiome, we used the Inverse Simpson α-diversity formula on the phylum level abundance counts (Shannon Diversity Index, Simpson, and Inverse Simpson measures are all highly correlated; see Table S9), as computed in Vegan (Jari Oksanen, F. Fuillaume Blanchet, Roeland Kindt, Pierre Legendre, Peter R. Minchin, R. B. O’Hara, Gavin L. Simpson, Peter Solymos, M. Henry, H. Stevens, Helene Wagner 2015). The Firmicutes-to-Bacteroidetes (F:B) ratio was calculated separately for gut and oral microbiomes of each child, and for oral microbiome of each mother, by taking the total count of the number of reads assigned to the Firmicutes phylum divided by the total count of the number of reads assigned to the Bacteroidetes phylum. Non-metric Multidimensional Scaling calculations were also performed using Vegan (Jari Oksanen, F. Fuillaume Blanchet, Roeland Kindt, Pierre Legendre, Peter R. Minchin, R. B. O’Hara, Gavin L. Simpson, Peter Solymos, M. Henry, H. Stevens, Helene Wagner 2015). See Fig. S6 for a detailed schematic of the computational workflow.

### Growth Curve Construction

At each time point when weight and height were collected (see above), we computed the ratio of weight to length (later referred to as growth index). These ratios were then analyzed longitudinally using tools from Functional Data Analysis (FDA) (Ramsay and Silverman 2005) alongside the *fda* package in *R.* In particular, individual growth curves were constructed using PACE (Yao et al. 2005), a procedure that pools information across subjects to more accurately assemble the curves (Fig. S2). The PACE software is freely available for R, and we used it with its default settings. After the curves were assembled, we represented them using 102 cubic spline functions with evenly spaced knots, so that subsequent FDA methods could be applied. Growth curves were also temporally aligned, using the *register.fd* function in R, before further analyses were conducted (Fig. 1B).

### Association between growth curves and microbiome summary measures

We assessed the association between growth curves and the α-diversity, as well as the F:B ratio, of gut and oral microbiomes fitting *Function-on-Scalar Linear Models (Kokoszka and Reimherr 2017)*. These were low-dimensional functional regressions (the growth curve response, i.e. the function, was regressed on a single scalar predictor: α-diversity or the F:B ratio), and were carried out in R using code we wrote based on (Kokoszka and Reimherr 2017). The outcome of these functional regressions are estimated regression coefficient curves, which we obtained using a penalized least squares approach imposing a penalty on the second derivative of the parameter. In each case, the smoothing parameter was set at 10,000, but results were robust against this choice (especially the p-values; see below). The additional smoothing enforced by the penalization was especially useful in terms of producing interpretable estimates. The shape of each estimated coefficient curve indicates how the relationship evolves along the time dimension, with amplitude (distance from zero) and sign (positive or negative) representing strength and direction. Significance was determined as described in (Choi and Reimherr 2017), based on three tests which employ different types of weighted quadratic forms. The first, denoted as L2, employs a simple L2 norm (squared integral) of the parameter estimate. The second, denoted as PCA, uses principal components to reduce the dimension of the parameter and then applies a Wald-type test. The last, denoted as Choi, incorporates a weighting scheme into the PCA test so that more principal components can be included, resulting in a test that is in between the PCA and L2 tests. Code is available at: (https://github.com/mreimherr/InsightMicrobiomeSimulation.git)

### Conditional Weight Gain score calculation

Age- and gender-specific weight-and length-for-age z-scores (WAZ and LAZ, respectively) were determined using the World Health Organization gender-specific child growth standards (The World Health Organization). Conditional weight gain (CWG) z-scores were then computed for each child using age-and gender-adjusted anthropometrics at birth and six months (Griffiths et al. 2009; Savage et al. 2016). Briefly, CWG z-scores were computed as standardized residuals from a linear regression of WAZ at six months on WAZ at birth, using LAZ and precise age at six months visit as covariates. The CWG z-score, therefore, represents the variability of child weight gain explained neither by length at birth and six months nor by gender. By construction, the CWG z-scores have mean 0 and standard deviation of 1. Moreover, in practice these scores are approximately normally distributed. Positive CWG z-scores indicate weight gain that is above the average weight gain, i.e. rapid weight gain.

### Testing potential co-factors

We tested a wide variety of maternal, health, and behavioural factors (gathered by the INSIGHT study; Paul et al. 2014) for effects on the children’s microbiomes, and the relationships of the microbiomes with children’s weight gain. Maternal gestational weight gain, diabetes during pregnancy, mode of delivery, and gender of the child were obtained from electronic medical health records. Maternal smoking during pregnancy, family income, child exposure to antibiotics or acid reducing medications, were obtained from maternal recall surveys.

First, considering these factors one at a time, we tested whether gut and oral microbiome diversity and (separately) F:B ratio differed between their categories using non-parametric Mann-Whitney U and Kruskal-Wallis tests implemented in R (Table S2).

We also obtained information on a child’s diet at two years as reported by parents using an Infant Food Frequency Questionnaire. This questionnaire included 121 food and drink items. Parents reported how often their child had each item in the past week (0, 1, 2-3, or 4-6 times per week and 1, 2, 3, 4-5, 6 or more times per day). These data were then distilled into a 10-item summary, each item with the corresponding number of weekly consumptions. The items included: sugar-sweetened beverages, milk, dairy (excluding milk), fruit, vegetables, vegetables excluding potatoes, snacks, sweets, meats, and fried foods. We looked at correlations between the consumption frequencies to see if any of the items could be removed (we used the R package *Rstats* and the graphical package *corrplot* (Wei 2013)). A correlation cut off of 0.7 was employed, eliminating the items milk and vegetables excluding potatoes. Consumption frequencies for the remaining eight food categories were then used as predictors in multiple linear regressions for the microbiome summary measures (four regressions in all, for diversity and F:B ratio in children’s oral and gut microbiomes; Table S3). This was performed using the *lm* function of the *Rstats* package.

Next, we used the *bestglm* package in R (McLeod and Xu 2014) to select the best subset of predictors for a multiple linear regression. This was performed again for four regressions (diversity and F:B ratio in children’s oral and gut microbiomes), this time considering seventeen covariates at our disposal, as listed in Tables S2 and S3. The covariates selected by *bestglm* were then used as predictors in restricted linear regression fits (results are reported in Table 1 for gut microbiomes’ α-diversity and F:B ratio, for which only diet-related covariates were retained; no covariates were retained by *bestglm* for the oral microbiome summary measures).

Given the prominent effects of diet-related covariates, especially on the gut microbiome (Tables S3 and Table 1), we also assessed their relationship with weight gain. Specifically, we ran a multiple functional regression for children growth curves (between birth and age two) against food consumption frequencies at age two (Table S4; we used the same eight food categories considered in Table S3). Like with the functional regressions for growth curves against microbiome summary measures (see above), statistical significance was determined with three different tests : L2, PCA, and Choi (Choi and Reimherr 2017).

Finally, we assessed whether diet-related covariates could modulate the relationship between weight gain and the gut microbiome. Specifically, we repeated the two functional regressions of children growth curves on gut α-diversity (Table S5) and gut F:B ratio (Table S6), in each case adding the diet-related covariates retained by *bestglm* (Table 1). Statistical significance was again determined with the L2, PCA, and Choi tests (Choi and Reimherr 2017).

### Identification of influential taxonomic groups

To mitigate sparseness and collinearity in our bacterial abundance data, we merged low-abundance or highly correlated genus-level abundance counts into abundances of taxonomic groups. We implemented a two-stage procedure which utilizes phylogenetic relationships – merging abundances only for neighboring nodes along the phylogenetic tree created from the Kraken taxonomic report tool in Galaxy (Wood and Salzberg 2014). First, moving upwards along the tree, we merge a node with its neighbor if its abundance is less than five counts in more than 90% of the samples in the data set. When such a merger occurs the counts from the two nodes are summed. Next, considering the merged abundances produced by the first stage and moving upwards along the tree, we merge neighboring nodes if their abundances show a correlation in excess of 0.7 across the samples in the data set. When such a merger occurs the counts are averaged. Notably, this procedure allowed us to tailor the level of resolution of our analyses to the data: branches of the phylogenetic tree where genera were scarcely observed or highly correlated were lumped together, while finer resolution was maintained for branches where genera were more abundant and diversified in their behavior across the samples. The procedure was applied, separately, to abundance data from the three microbiome types (child gut, child oral and mother oral). Table S7 contains a complete list of all taxonomic groups obtained in each microbiome type.

To identify taxonomic groups with the strongest associations with weight gain, we considered the merged taxonomic group abundances as scalar predictors in functional regressions for growth curves using FLAME (*Functional Linear Adaptive Mixed Estimation*; Parodi and Reimherr 2017). Separately, we considered the merged taxonomic group abundances as features in Linear Discriminant Analysis for rapid vs. non-rapid weight gain (based on CWG scores) using LEfSe (Linear Discriminant Analysis Effect Size; Segata et al. 2011) as implemented in Galaxy (Blankenberg et al. 2010b). These were high-dimensional analyses (each comprised a number of predictors corresponding to the number of taxonomic groups found in a given microbiome type). The FLAME functional regressions were carried out using methods and R code from (Parodi and Reimherr 2017). Estimates and p-values were computed using standard methods from FDA (see github repository link below for code) as described in (<u>Ramsay and Silverman 2005</u>). FLAME simultaneously selects important predictors and captures their effects as estimated regression coefficient curves. It can be thought of as a generalization of the adaptive LASSO (Zou 2006) to functional/longitudinal outcomes. In particular, a Sobolev kernel was used in the penalty to produce smooth estimates while also carrying out variable selection. The tuning parameters were chosen via cross validation, as discussed in (Zou 2006; Parodi and Reimherr 2017). Code and examples for carrying out the two-stage abundance merger procedure and all of the FDA methods utilized here can be found at <https://github.com/mreimherr/InsightMicrobiomeSimulation.git>. Standalone code for FLAME is available at: <http://personal.psu.edu/~mlr36/codes.html>.

### Data Sharing

Raw microbiome reads were deposited into dbGaP Study number XXX. All code used is either already public or available at GitHub. 16S rRNA gene analysis pipeline tools and pipeline are available in the Galaxy platform (usegalaxy.org). The three relevant Galaxy workflows are https://usegalaxy.org/u/sjcarnahancraig/w/16s-qc-3, https://usegalaxy.org/u/sjcarnahancraig/w/kraken-classification, https://usegalaxy.org/u/sjcarnahancraig/w/vegan-rarefacation--alph-diversity.

### Ethics statement

This study was approved by Penn State University Institutional Review Board (PRAMS034493EP). Informed consent was received from mothers prior to collection of biological samples and phenotypic, demographic, health, and diet information.

## ACKNOWLEDGEMENTS

We would like to thank B. Higgins, A. Shelly, P. Carper, J. Beiler, N. Verdiglione, L. Hess, and the Penn State Genomics Core Facility, University Park, PA for their assistance. This project is supported by grants R01DK088244 and R01DK99364 from the National Institute of Diabetes and Digestive and Kidney Diseases (NIDDK). This project is also supported by the funds from the Eberly College of Sciences at PSU, by the Penn State Institute of Cyberscience, by the National Center for Research Resources and the National Center for Advancing Translational Sciences, National Institutes of Health (through Grant UL1TR000127). The content is solely the responsibility of the authors and does not necessarily represent the official views of the NIH. Additionally, this project is funded, in part, under a grant with the Pennsylvania Department of Health using Tobacco Settlement and CURE funds. The Department specifically disclaims responsibility for any analyses, interpretations, or conclusions.

